# Physiologic flow-conditioning limits vascular dysfunction in engineered human capillaries

**DOI:** 10.1101/2021.03.03.433247

**Authors:** Kristina Haase, Filippo Piatti, Minerva Marcano, Yoojin Shin, Roberta Visone, Alberto Redaelli, Marco Rasponi, Roger D. Kamm

## Abstract

Hemodynamics play a central role in the health and disease of the coronary and peripheral vascular systems. Vessel-lining endothelial cells are known mechanosensors, responding to disturbances in flow – with mechanosensitivity hypothesized to change in response to metabolic demands. The health of our smallest microvessels have been lauded as a prognostic marker for cardiovascular health. Yet, despite numerous animal models, studying these small vessels has proved difficult. Microfluidic technologies have allowed a number of 3D vascular models to be developed and used to investigate human vessels. Here, two such systems are employed for examining 1) interstitial flow effects on neo-vessel formation, and 2) the effects of flow-conditioning on vascular remodelling following sustained static culture. Interstitial flow is shown to enhance early vessel formation via significant remodeling of vessels and interconnected tight junctions of the endothelium. In formed vessels, continuous flow maintains a stable vascular diameter and causes significant remodeling, contrasting the continued anti-angiogenic decline of statically cultured vessels. This study is the first to couple complex 3D computational flow distributions and microvessel remodeling from microvessels grown on-chip (exposed to flow or no-flow conditions). Flow-conditioned vessels (WSS < 1Pa for 30 micron vessels) increase endothelial barrier function, result in significant changes in gene expression and reduce reactive oxygen species and anti-angiogenic cytokines. Taken together, these results demonstrate microvessel mechanosensitivity to flow-conditioning, which limits deleterious vessel regression *in vitro*, and could have implications for future modeling of reperfusion/no-flow conditions.

## Introduction

The importance of hemodynamics in the development and maintenance of cardiovascular health is unparalleled. During formation of neo-vessels, mechanosensitivity of endothelial cells is reduced to allow for their rapid reorganization [1], whereas fully-formed vascular networks are highly sensitive to minor changes in flow. In both large and small vessels, altered flow can arise from vessel occlusion, constriction, reduced cardiac output and/or vascular damage. Vessels adapt by releasing nitric oxide (a vasodilator), thereby regulating inflammation and proliferation of parenchymal cells. However, overly-stressed flow regimes (outside a tolerance threshold), results in reactive oxygen species (ROS) and other factors, including endothelin, to be over-actively released from the endothelium leading to vessel constriction, tissue inflammation and hypoxia [2]. For instance, re-introduction of blood flow to ischemic tissues (an extreme change in flow conditions) can lead to significant damage of vessels, termed ischemic reperfusion injury (IRI) [3]. Reperfusion of microvessels is often associated with acute infiltration and accumulation of leukocytes, vasoconstriction, and endothelial damage. Thus, investigating flow-conditioning and the threshold of microvessel flow-sensitivity demands further study.

Humanized *in vitro* systems provide an opportunity to understand complex microvessel hemodynamics in a highly controlled and simplified setting. Assessing microvascular function *in vivo* has historically proven difficult; however, recent acoustic and optical imaging techniques now infer the health of microvessels in response to vasodilators/constrictors [4, 5]. Animal models have also provided a wealth of information regarding vascular function; yet, these models fail to address the role of microvessels [6], and cannot assess the vascular response to altered flow systematically. Microfluidic models have proven particularly useful for controlling fluid flow in 3D vessels on-chip (reviewed in [7]). For example, angiogenesis assays have demonstrated the effects of interstitial flow (IF) over a variety of magnitudes and in relation to morphogen gradients including vascular endothelial growth factor (VEGF) [8–11]. Within the physiologic range (0.1-10 μm/s), IF has been shown to be the sole director of angiogenesis, as biochemical gradients are completely lost in several hours [11]. Most of the work reported thus far has focused on angiogenic effects, and only recently has IF been shown to have a significant impact on the growth of newly formed vessels [9]. One aim herein is to address this gap in knowledge by characterizing the effect of IF on neo-vessel morphology and connectivity.

Besides IF, microfluidic models have also been used to examine luminal flow disturbances, which can lead to endothelial dysfunction. Measurements *in vivo* for wall shear stress (WSS) in human capillaries range from ^~^0.2-10 Pa (in conjunctival vessels) [12], with values of WSS being inversely proportional to vessel diameter. In response to stressed flow conditions (beyond this range), abnormal vascular reactivity, increased permeability to solutes and expression of adhesion and chemotactic molecules, recruitment of leukocytes, changes in endothelial and stromal cell viability, and increased coagulation and thrombosis (von Willebrand Factor (vWF) and tissue factor), have all been reported [13]. *In vitro* models have shown these deleterious effects, for instance, by culturing primary human endothelial cells under disturbed flow regimes; however, the majority of this insight derives from planar cultures [14, 15]. More recently, 3D patterned vessels (straight and tortuous) have been used to demonstrate WSS effects from a variety of magnitudes, with vWF release occurring at a minimum of 0.3 Pa [16], and others demonstrating the requirement for NOTCH1 signalling to maintain endothelial barrier function in response to flow [17]. Most computational vascular models to-date have employed simplified geometries from physical reconstructions of *in vivo* or *in vitro* vessels [18]. Straight and spiraled vessels (of known dimensions) demonstrate laminar and complex WSS gradients under various flow regimes, respectively [19]. Despite these advances, an approach to model fluid dynamics in complex heterogenous human microvessels has been lacking.

Coupling self-assembled vessels with computational modeling, this work aims to examine microvascular hemodynamics in response to flow-conditioning. Microfluidic vascular models are employed to 1) demonstrate the impact of interstitial flow on neo-vessel formation, and 2) provide insight into the global effect of continuous flow-conditioning on the remodeling behaviour of heterogeneous human microvessels. Early formation and morphologic changes result from IF, which caused significant remodeling of tight junctions in neo-vessels. By culturing vessels under continuous flow, microvascular health was sustained through remodeling. Computational fluid dynamics revealed heterogeneous flow distributions with wall shear stresses and shear rates similar to those expected for precapillary venules (< 1Pa and < 500s^−1^) – with higher magnitudes in flow-conditioned than in prolonged static cultured vessels. Long-term flow-conditioning marginally altered endothelial barrier properties, significantly altered gene expression and reduced expression of ROS and inflammatory factors. Taken together, a highly tuned mechanosensitivity of human microvessels is demonstrated, which causes remodelling and reduced ROS in response to low wall shear stresses following static culture, suggesting that pre-conditioning microvessels may alleviate cell death and tissue ischemia.

## Results

### Interstitial flow enhances vessel growth and remodelling

Fluid dynamics play an important role in the formation and maintenance of the microcirculation; however, only little is known about the effect of interstitial flow on neo-vessel formation [9]. Our well-established methods for culturing *in vitro* vessels from human primary cells are primed for investigating the effect of IF on the process of vasculogenesis. First, microvessels were cultured using human umbilical vein endothelial cells (HUVECs) tagged with an RFP cytoplasmic marker and human lung fibroblasts (HLFs) seeded in a 5:1 ratio in a fibrin hydrogel, as done previously [20, 21]. Commercially available microfluidic devices (AIM Chips) were employed to facilitate the establishment of a pressure gradient across the hydrogel (using attachments and syringes as shown in Figure 1A). IF was re-established 3x daily (every 4 hours – to account for the loss of pressure, see Figure S1A) in each chip by ensuring a ^~^2.5mm H_2_O pressure difference across the gel. Fluorescence recovery after photobleaching (FRAP) was used in cell-free gels to estimate the mean IF initial velocity as ^~^5 μm/s (extrapolated from linear fit in Figure S1 B,C), as done previously [10].

**Figure 1:**
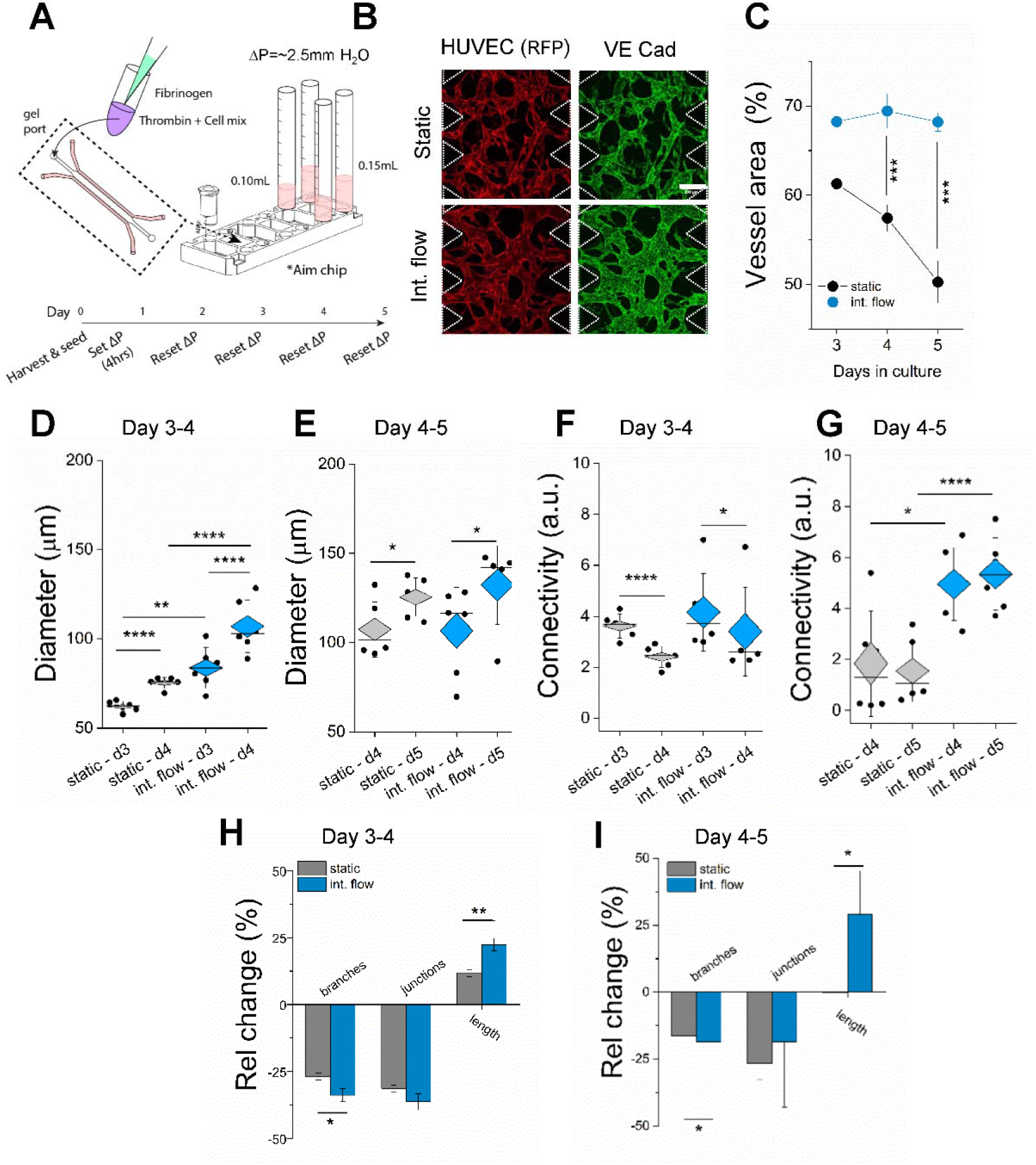
Interstitial flow promotes early vessel growth and connectivity. A) Schematic diagram of set-up for interstitial flow and timeline using AimChip and affixed 1mL syringes (shown with media volume and set pressure difference). B) Fixed image of vessels exposed to either static or interstitial flow conditions (day 4 shown). Scale bar is 200μm. C) Mean vessel area coverage is shown for 3 days in culture. n= 6 devices were analyzed for days 3 and 5, n=12 for day 4. One dot is equivalent to 3 measurements made per device. Vessel morphologic parameters are shown for days 3-4 (D,F,H) and days 4-5 (E,G,I). D) and E) show the mean effective vessel diameters. F) and G) report connectivity as a ratio of junctions/endpoints. H) and I) show relative changes in the indicated parameters, compared to the same device from the prior day. Significance is indicated by unpaired t-tests between static and flow and paired t-tests between the same condition, with *P<0.05, **P<0.01, ***P<0.001, ***P<0.0001.

To track vessel growth, cells were imaged using confocal microscopy (volumetric imaging) starting at day 3, at which time nascent vessels began to form. Vessel morphology was noticeably altered by IF treatment, as shown by fixed images and staining (Figure 1B). Endothelial migration was observed in the media channels for IF-condition devices with vessels appearing to align in the flow direction (Figure S2A). While increased proliferation (ki67+ cells) was not detected, overall cell count was higher for IF conditions (Figure S2 B-D). For separate experiments, live images were captured on days 3 and 4, as well as days 4 and 5, respectively, in order to track morphologic change in response to static and IF culture conditions. Several parameters (i.e. vessel projected area, branches, junctions, vessel length, diameter) were analyzed by post-processing the images in ImageJ (Figure 1C-I), as done previously [20]. Quantification of these parameters led to a confirmation that IF has a significant effect on neo-vessel morphology. Vessels exposed to IF cover significantly more area over time (Figure 1C), and by day 3 present significantly increased effective mean diameter values, in comparison to vessels grown statically (Figure 1D). Drastic differences in mean diameter is time-dependent, and is less extensive by day 4-5, as seen by comparing IF and static cultures in Figure 1E. Interestingly, connectivity of the vessels (defined as a ratio of junctions/endpoints) decreases early on, between days 3 to 4, under both static and flow conditions (Figure 1F). Presumably, this occurs due to initial pruning of the neo-formed vessels, prior to any further branching – we observed a similar trend previously in static cultures [20]. By day 4, connectivity of those vessels treated with IF is significantly increased, suggesting further vessel remodeling (Figure 1G).

Changes in morphologic parameters were calculated across days 3-4 and days 4-5 of culture (Figure 1 H, I). Number of branches, junctions and average branch length were measured for a specific device on consecutive days, thus quantifying relative changes – which reduces variance in initial seeding and/or vessel distributions. Significant change was apparent in branch number and length between vessels grown under static or IF conditions across both days. In the case of IF, significant pruning and remodelling was evident based on the relative reduction in branches and increase in branch length, respectively. Staining demonstrated a reduction in tight junction stability due to IF (indicated by reduced ZO-1 intensity, Figure S2 E-F). This loss of junction stability may be due to the intermittent application of IF and may be transient, as similar temporary decreases in adherens junctions proteins have been seen previously in the short-term following application of flow on endothelial cells [22].

Overall, IF initiated early vessel formation and remodelling in comparison to statically cultured vessels. Vessel formation occurred quite rapidly (several days) in these micro-scaled chips, with connections appearing in the media channel. Thus, maintaining true IF over time in this setup was not possible, since the gradient transitioned to combined IF and vascular luminal flow. Given that open lumen appeared by day 4 (Figure S2A), flow is expected to be mostly intraluminal at that time.

### Long-term (24hr) continuous flow induces significant vascular remodeling

To examine the effects of continuous flow on vascular remodeling, macro-scale PDMS devices (gel region of 2.5mm length x 3mm width x 1mm height) were used for culturing vessels (Figure 2A). In these larger devices (fabricated in-house), vessels formed over the course of one week, and are fully perfusable by day 7, as we have shown previously in similar but slightly smaller devices [20, 21]. Vessels were cultured under static conditions for 7 days prior to connecting a large media reservoir, which was then used to generate a pressure gradient across the gel (Figure S3 A,B). The reservoir was integrated with a controlled solenoid driven air pump, to maintain fluid recirculation at a constant pressure gradient of ^~^5mm H_2_O (^~^50Pa) across the gel and vessels. Microvessels were imaged at day 7 (prior to flow), 8 (24 hours of flow), and 9 (48 hours of flow) using confocal microscopy (examples at 48 hours are shown in Figure 2B). Flow was temporarily suspended to image the devices, followed by a subsequent media top-up (to account for slight evaporation) and return to continuous flow for a second consecutive 24-hour conditioning period.

**Figure 2:**
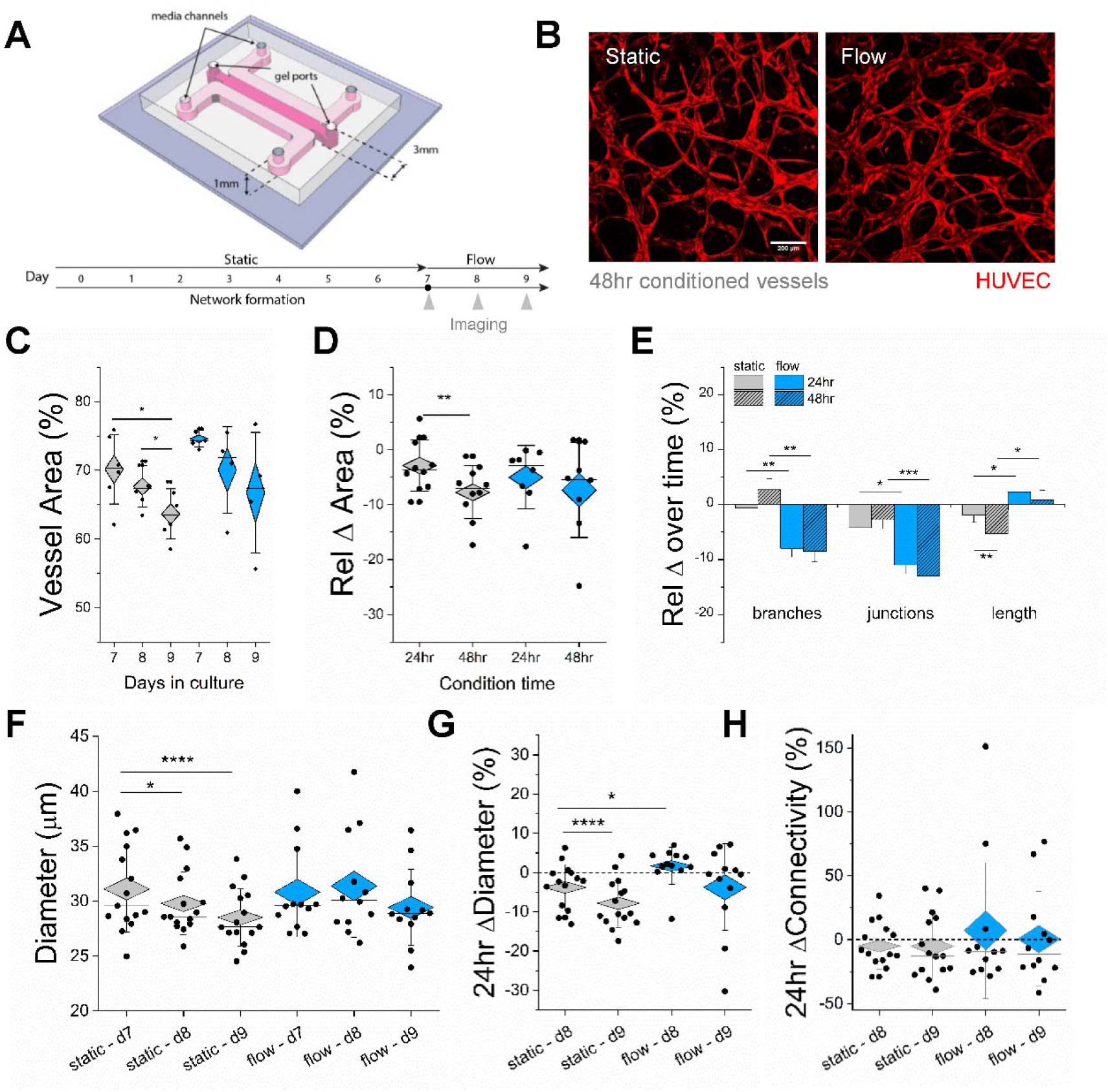
Continuous flow applied to formed vessels promotes sustained vessel structures. A) Cartoon of microfluidic device and timeline for vessel formation and application of flow/static conditioning. B) Examples of vessels after 48hrs of either static or flow-conditioned vessels (day 9). C) Quantification of vessel area coverage for both static (grey) and flow (blue) conditions from day 7 to 9. D) Relative change in vessel area between 24 and 48hrs of the two conditions – static (grey) and flow (blue). E) Relative change in number of branches, junctions, and in mean branch length, as seen over 24 and 48hrs of either static or flow conditions. F) Mean diameter as shown for samples measured over consecutive days (7-9) for each condition. G) Relative change in diameter corresponding to (F) over 24 and 48hrs. H) Relative change in connectivity (measured as a ratio of vessel junctions/endpoints) over time. Each dot represents a single device and mean of 3 measures made per device. Significance is indicated by unpaired t-tests between static and flow and paired t-tests between the same condition, with *P<0.05, **P<0.01, ***P<0.001, ***P<0.0001.

From the images collected, it was possible to measure changes in vessel growth and morphology in response to static or flow-conditioning (as with IF). First, vessel area coverage was measured and shown to decrease for both conditions over time (days 8 and 9); however, the decrease was only significant for those in static culture, unlike those exposed to flow (Figure 2C). This decrease in vessel growth was corroborated by measuring the relative change in area for each device (Figure 2D). Accordingly, vessels grown under static conditions undergo significant thinning in comparison to those pre-conditioned by flow. Next, other relevant vessel remodeling parameters were compared between days 7-8 and days 8-9 for static and flow conditions (Figure 2E). Interestingly, relative changes in the number of branches and junctions were significantly decreased for vessels conditioned by continuous flow (in both 24- and 48-hour periods), suggesting that vessels exposed to flow undergo significant remodeling. In line with remodeling in the absence of vessel thinning, flow-conditioned vessels also result in significant increases in mean vessel length, in comparison to those cultured statically.

Vessels conditioned by continuous flow exhibited less vessel rarefaction. This is also demonstrated by measuring mean vessel effective diameter (Figure 2F,G). Diameter was significantly reduced under static culture conditions, as shown by absolute and relative measurements, respectively. In the case of flow, the relative change in diameter increases for the first 24hrs of treatment. Neither condition led to any significant increase or loss of vessel connectivity in 48 hours (Figure 2H). However, longer cultures with flow (96 hrs) eventually result in vessel rarefaction (Figure S3 C). To approximate the fluid flow velocity in these microvessels, 10 μm fluorescent beads were perfused at two pressure gradients demonstrating an expected increase in mean velocity in response to increased pressure, and a heterogeneous flow distribution in the vessels. Velocities measured by particle image velocimetry (PIV) tracing, as expected, (Figure S3D-F) demonstrated an increase in velocity for each of n=3 devices upon increasing the pressure difference. Velocities for 5mm H_2_O pressure difference used for flow-conditioning experiments are expected to be ^~^0.24 mm/s (extrapolated from linear fit), but could not be tracked due to limitations in detector frame rate. Taken together, it appears that continuous flow-conditioning can offset the early onset of vessel rarefaction, but cannot altogether avoid the limitations of longterm culture *in vitro*, under the conditions set in these experiments.

### Computational fluid dynamics of vessels highlights complex flow patterns

Considering that our *in vitro* vessels cannot be equated to a series of simplified tubes and PIV only approximates fluid flow, we aimed to generate a computational fluid dynamics (CFD) simulation capable of providing a clear picture of flow in our microvessels. First, large-scale regions of the vessels were imaged using confocal microscopy on day 9, from vessels cultured under both static and pre-conditioned flow (n=3 each). A video demonstrating one representative 3D region is shown in Video S1. These images were stitched and a threshold was applied in Matlab in order to generate an iso-surface to define the vessel walls, as in [23]. Numerical modeling was performed using ANSYS ICEM CFD, the full details of which can be found in the Methods section. Briefly, laminar flow and a no-slip boundary was simulated at the vessel wall. The fluid (culture media) was considered Newtonian, having a dynamic viscosity of 9.4×10^−4^Kg·m^−1^·s^−1^ and density of 998.2 kg·m^−3^ at 37°C (values previously reported for DMEM +10%FBS [24]). Fluid flow was then simulated with a similar pressure drop across the gel as in the *in vitro* experiment (see Figure S4). An example of a simulation of one unstitched region of experimentally-derived vessels is shown to demonstrate the complex fluid flow velocity and WSS in greater detail, as seen in Figures 3A and 3B, respectively.

**Figure 3:**
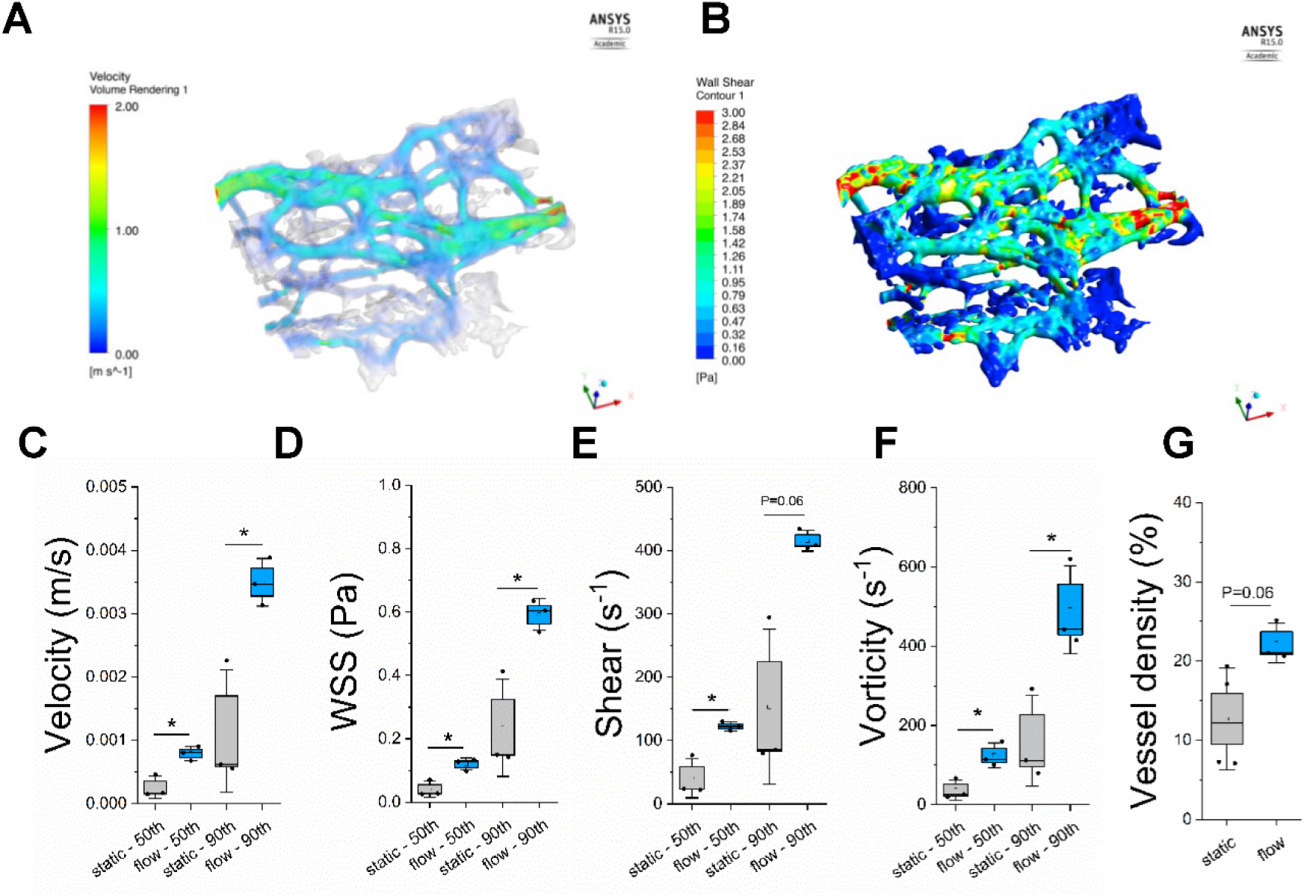
Computational characterization of flow in conditioned *in vitro* vessels.A) Example of one region of reconstructed vessels where CFD is performed to indicate fluid velocities and corresponding B) WSS measurements. C-G) Various fluid flow parameters were measured for 3D reconstructions of samples cultured under static or flow conditions (day 7-9). E) Mean fluid velocity is shown for static and flow conditions, as well as corresponding F) WSS, G) shear rate, H) vorticity (curl of flow velocity), and I) vessel density (% volume). Shown are box plots with outer box SE and error bars as SD. Both the means of the 50^th^ and 90^th^ percentiles of all simulated measurements are shown. Significance is indicated by unpaired t-tests between static and flow, with *P<0.05.

Several parameters were computed from the simulations, including: mean fluid flow velocity, WSS, vorticity (curl of flow velocity), and % volume vessel density (Figure 3C-G). Both the 50^th^ and 90^th^ percentiles of the mean computed values were reported here in order to compare between simulations from reconstructions of static- and flow-conditioned vessels. Given the same simulated pressure drop across the vessels, the thinning of statically cultured vessels led to distinct differences in flow patterning (fewer perfusable segments) in comparison to flow-conditioned vessels (Figure S4). The vessels reconstructed from flow-conditioned samples demonstrated a more homogenous fluid flow distribution (more perfused segments and more uniform fluid flow characteristics). This distribution can be demonstrated by a computed threshold velocity (V_t_), here defined as Vmax/10. On average, 22.5 ± 1.9 % of the velocities were below this threshold for flow-conditioned vessels, whereas 35.3 ± 15.3 % were lower than this threshold for static vessels. Flow-conditioning alters the morphology of the vessels (and maintains diameter), and thus led to significant increases in all of the aforementioned parameters. Overall, for both static- and flow-conditioned samples, velocity measurements at a pressure difference of 5 mm H_2_O are within the range expected (several mm/s) for pre-capillaries [25]. Moreover, the WSS (mean of 0.5 Pa for flow or 5 dyne/cm^2^) is also within range of reports for human conjunctival capillaries [12]. Overall, low levels of luminal fluid flow induced changes in vessel geometry and density, leading to more perfused regions and increasingly even WSS distributions in our vessels following static culture.

### Flow-conditioning alters vessel behaviour function and gene expression

Our results have clearly shown flow-induced changes in vascular morphology occurring over 1-2 days. However, we also aimed to demonstrate the responsiveness of applied flow in the vessels. Therefore, we performed an experiment to detect the sensitivity to acutely applied flow, by perfusion of vessels at a pressure drop of 2 mm H_2_O at day 7 with 4,5-Diaminofluorescein (DAF-2) – a fluorescent indicator of nitric oxide (NO). After incubation with the dye, vessels were imaged under static conditions (8 mins) and then flow was induced for a consecutive 8 min period (by addition of media to incorporated syringes, Figure 4A). Time-lapse imaging was performed in order to capture any increase in fluorescence in the vessels, as was stimulated by flow, as evidenced in Figure 4B. A progressive increase in NO was shown for vessels stimulated by flow, in comparison to static vessels where no change was detected over time.

**Figure 4:**
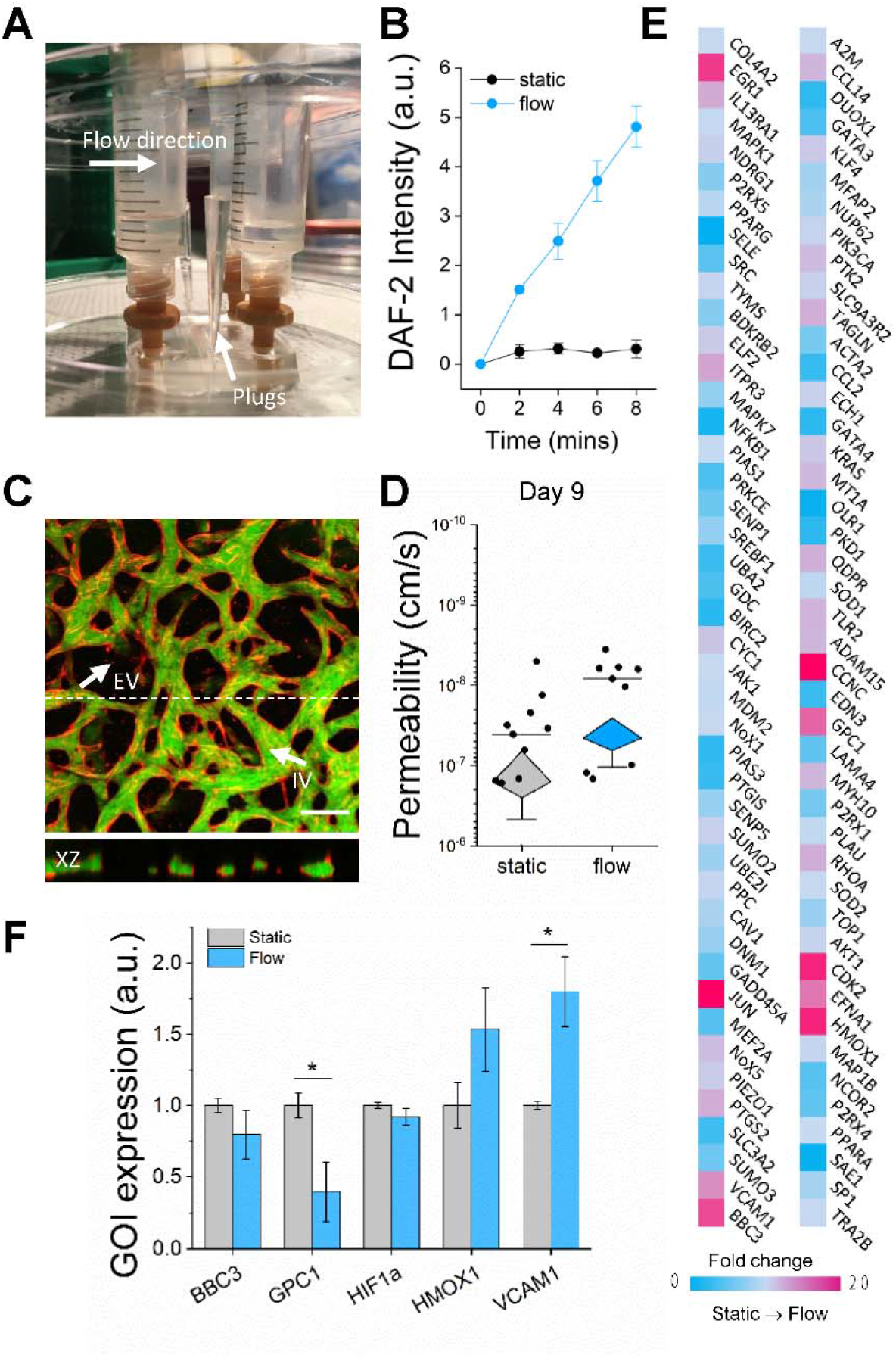
In vitro vessels are responsive to short and long-term flow-conditioning. A) Image of setup for short-term flow induction. Plugs are placed in the gel ports to sustain the pressure gradient across the gel. B) DAF-2 (normalized) intensity is shown following immediate continuous flow versus no flow over 8 minutes. C) Confocal z-stack of vessels (HUVEC – red) perfused with 70kDa FITC (green) dextran after 48hrs of continuous flow (day 9). Arrows indicate intravascular (IV) and extravascular (EV) regions. Scale bar is 200 microns. D) Permeability measurements are shown for several experiments made on day 9 (following 48hrs of continuous flow or static growth conditions). Shown are mean values for individual devices (2-3 measurements per device). E) Plot of fold change between static and flow conditions for angiogenesis gene query array. F) Genes of interest shown as normalized fold-change to static conditions. All box plots are the height of SE and tails SD. Significance is indicated by unpaired t-tests between static and flow, with *P<0.05, **P<0.01.

Considering that the microvessels in this system are clearly mechanosensitive, we next aimed to look at the long-term (48 hour) effects of flow-conditioning on the endothelial barrier, as flow is known to alter this vascular property (with laminar WSS reducing permeability) [17]. As we have done previously [20], we employed FITC dextran (70kDa) to measure changes in permeability (P_e_) that might follow 48 hours of flow-conditioning. Flow was suspended, and a pressure gradient was introduced across the gel to perfuse the fluorescent dextran through the vessels, following which an equal volume of media (100μL) was added to the opposite media channel (to stop convective flow). Confocal z-stacks were captured over time (5 min intervals) and flux of dextran from intravascular to extravascular regions (see Figure 4C) was later measured by post-processing the images in ImageJ. Flow-conditioned vessels result in increased barrier function to solutes (mean values for P_e_ are 4.6×10^−8^ cm/s versus 1.6×10^−7^ cm/s for flow and static, respectively, Figure 4D), as expected from earlier studies on larger *in vitro* vessels [17, 26]. However, the difference is non-significant (P=0.293), since despite several independent biological repeats, variance was high between individual devices measured.

Next, to demonstrate the impact of flow-conditioning, we measured changes in vessel-stromal gene expression by pooling several samples together (n=3 each for flow and static). An angiogenesis-specific array was used to highlight genetic changes that occur following 48 hours of applied flow-conditioning. A number of genes were altered and resulted in large fold-changes (Figure 4E). Many of these genes were down-regulated, including those related to angiogenesis and proliferation (GPC1 and EFNA1), genes related to the cell cycle (CCNC and CDK2), mitogenesis (EGR1), and an AP-1 transcription factor subunit (JUN). Many of these genes have been previously demonstrated to be altered by laminar shear stress in HUVEC, which has been shown to contribute to growth inhibition [27]. We examined several specific genes of interest (GOI) across pooled samples from several individual experiments. RT-qPCR revealed significant decreases confirmed in GPC1 and increased VCAM1 expression, as expected in endothelial activation (Figure 4F). GPC1 is a heparan sulfate core protein that has been shown to regulate endothelial-cell NO synthase release – acting as a mechanotransducer to shear flow [28]. The response to shear flow (1.5 Pa) has been shown to result in transient clustering of GPC1 (reduction in expression) over short durations [29]. Moreover, increased levels of VCAM1 expression has been previously associated with low shear stress (0.2-0.4 Pa), as shown in endothelial monolayers [30].

### Anti-inflammatory effects are attributed to flow-conditioning

Since low flow-conditioning promotes NO release, we endeavoured to measure changes in hypoxia (HYP) and reactive oxygen species (ROS) using fluorescent stress-indicator dyes (ROS-ID). At day 9, no measured differences were detected in HYP between static- or flow-conditioned vessels (Figure 5A). This could be partially due to the fact that flow had to be stopped and the vessels incubated for a ½hr duration prior to measurement, or due to the nature of oxygen exchange through the PDMS device. On the other hand, flow-conditioned vessels did demonstrate a significant decrease in ROS, compared to statically cultured vessels (Figure 5B). Since ROS is associated with inflammation, we also examined whether there was any change in apoptosis using a fluorescent indicator of caspase 3/7; however, no change was detected (Figure 5C). Finally, two common cytokines strongly associated with vessel growth (angiopoietins (Ang) 1 and 2) were quantified in supernatants collected from vessels following 24 hrs of flow by ELISA. Ang1 is typically produced by smooth muscle cells and is associated with vessel stabilization and increased barrier function, while Ang2 is associated with vessel destabilization/remodelling [31]. While Ang1 was increased for flow (P=0.149), a ratio of these two commonly examined cytokines (Ang2/Ang1) demonstrates an overall decrease for flow-conditioned vessels (Figure 5D). This ratio is often used as an indicator of inflammation and shows positive correlation with disease severity [32, 33]. Taken together, these results suggest that low levels of shear flow-conditioning maintains health of our microvessels.

**Figure 5:**
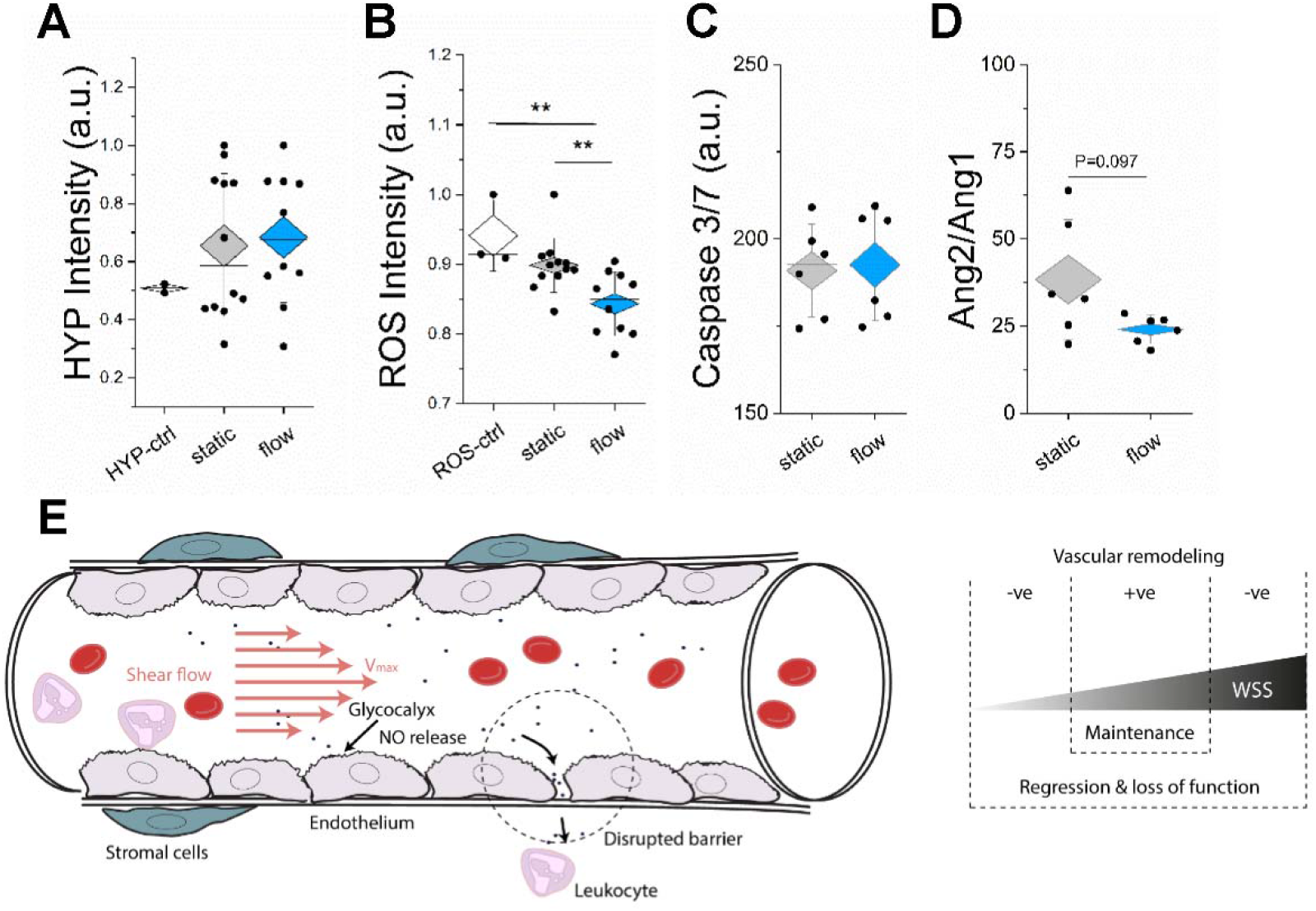
Inflammation is reduced in low flow-conditioned microvessels. A) Hypoxia measurements made following 48hrs of flow or static conditions. B) Reactive oxygen species (ROS) measurements made following 48hrs static or flow conditions (in comparison to a control). C) Caspase 3/7 measurements (mean intensity) shown for both static and flow conditions. D) Ratio of angiopoietin 2/1 (measured by ELISA) shown for static and flow conditions. E) Schematic representation of the effects of a flow on vascular remodelling and function. It is hypothesized that for the microvessels of our size, the fluid flow promoted vascular maintenance, whereas continued static culture led to regression and loss of function (leakier vessels and increased ROS).

Overall, our results have shown the positive effect of interstitial and low shear flow-conditioning on *in vitro* microvessels. In particular, interstitial flow promotes early vasculogenesis and vessel connectivity. Moreover, following *de novo* vessel formation, continuously applied low levels of intravascular shear flow activates endothelial cells, but reduces vessel thinning and promotes anti-inflammatory effects. For the mean vessel diameters in these microvessels (^~^30 microns), the low shear stress magnitudes (^~^0.6 Pa) applied herein are within the threshold range of having a protective effect (anti-inflammatory). We propose that high and low magnitude flow-conditioning outside of this threshold results in regressive vascular remodeling – which contributes to a pro-inflammatory response (Figure 5E). We aim to examine the limits of this threshold – i.e. the sensitivity of our microvessels to this interplay between fluid flow properties and morphologic homeostasis further in future work.

## Discussion

Hemodynamics play a critical role in development, homeostasis and dysregulation of microvessels. However, the strong impact of disturbed flow, as occurs in vessel reperfusion, on morphogenesis of vessels is difficult to observe *in vivo*, making *in vitro* vessels a logical choice for its study. Herein, 3D human vessels were cultured on-chip to examine the resultant morphologic and functional changes due to flow-conditioning. First, we examined interstitial flow effects on early vasculogenesis, which revealed the mechano-responsiveness of nascent vessels.

Alignment of vessels in the direction of flow and early network formation and remodelling seen in our vasculogenesis experiments corroborates those results that were performed in a similar physiologic range of flow in earlier bioreactor systems [34], and is close to those seen in animal studies [35]. IF is known to promote sprouting angiogenesis and formation of 3D vessels in a magnitude-dependent manner [8, 36], acting concurrently with morphogen gradients to direct endothelial cell migration [11]. Previous work has shown that sensitivity to flow in early vessel formation is largely reliant on endothelial-derived Wnt ligands, which play a crucial role in endothelial polarization [37]. Moreover, microfluidic models with precisely controlled pressure gradients revealed that a vasculogenic response is dependent on the Peclet number (ratio of convective to diffusive flow) and that a range of insensitivity (0.1<P_e_<10) exists [38]. Using our mean IF flow velocity (V = 5 μm/s), a characteristic length of the AimChip (L = 1.3mm) and an estimate for diffusivity of a 70 kDa solute (similar to MW of Ang-2) in the gel (D = 250 μm^2^/s based on our earlier work [21]) we calculated a Peclet number P_e_=VL/D=26, which is much higher than the reported value needed to initiate a vasculogenic response. Importantly, we also demonstrated that IF results in remodeling behaviour following neo-vessel formation and transiently affects endothelial tight junctions. Mechanotransduction of IF has been previously shown to direct the migratory response of tumor cells embedded in hydrogels [39] as well as directing angiogenesis [40] through basal-apical gradients in matrix adhesion. Thus, when IF is present and detectable, remodeling behaviour is expected due to force-balancing by the vessels and stroma.

The responsiveness of endothelial cells to changes in shear flow has been shown in a variety of *in vitro* [15] and *in vivo* models [1]. Endothelial cells align with the direction of flow by polarized migration, thus being attracted towards well-perfused regions, resulting in regressive remodelling of poorly perfused vessels [1]. Using a macroscale fluidic device, we aimed to illustrate the effect of flow-conditioning on the morphogenesis and functionality of fully-formed *in vitro* microvessels. Following static culture of perfusable vasculature, flow-conditioning in our microvessels resulted in sustained vessel diameters (in comparison to the narrowing of ischemic-like static vessels), which has been shown earlier in WSS-induced remodeling in animal studies (reviewed in [41]). On average, the permeability to solutes was decreased (improved barrier function) due to flow-conditioning; however, static vessels also resulted in relatively stable (non-leaky) vessels with similar values to those reported in animal studies (values on the order of 10^−7^ tabled in [42] for various tissues of murine models). A number of genes related to the stress response, remodelling, and proliferation were upregulated in response to flow in the short (48 hour) window of treatment. Moreover, flow-conditioning resulted in an antiinflammatory response – demonstrating reduced ROS and a reduction in the Ang2/Ang1 ratio, which has been used as a prognostic indicator of disease [33]. Importantly, our results demonstrated that low levels (<1 Pa) of flow-conditioning can majorly impact microvascular health in comparison to continued static/ischemic-like culture.

CFD models have been used to elucidate complex fluid patterns inside vessels reconstructed from anatomic or microscopy-based images. Some of these models include investigation of pulsatile flow in vessels representative of stenosis [43], volumetric flow rates after carotid artery stenosis [44], as well as the interactions and contributions of deformable cells (including RBCs or circulating tumor cells) to disturbed flow in microchannels [45, 46]. Here, to assess the effect of flow-conditioning on developing flow patterns, we coupled a CFD model to our experiments using several reconstructed vascular geometries from no-flow and flow-conditioned samples. Strikingly, the flow patterns that developed in our vessels revealed heterogeneous yet preferential flow regions, with flow-conditioning resulting in more perfused segments overall (an increased vessel density). Thus, our CFD modeling supports the hypothesis that regression likely occurs in non-perfused segments, by inferring that a reduction in flow (below a threshold) coincides with reduced vessel density, as seen experimentally. Parameters including velocity and WSS were assessed by making a number of assumptions to simulate fluid flow in our microvessels. For example, the fluid is considered Newtonian (an acceptable assumption for cell culture media) and vessel walls are considered to be inelastic with no remodeling. Future work will focus on incorporating the active remodeling of vessels, and complex 2-phase effects caused by the inclusion of red blood cells into these simulations, which will undoubtedly alter the flow velocity profiles and shear rates within the vessels, as has also been recently shown by capillary dilation *in situ* [47]. The WSS and shear rates at the 90^th^ percentile distribution of our simulation demonstrate values similar to those reported for post-capillary venules (2 Pa or 20 dynes/cm^2^), as measured previously *in vivo* for similarly sized vessels [48]. We note here that the simulations were performed at 100 Pa, but due to linearity in results, we can infer the values reported at the 50 Pa pressure difference in our experiments. Low shear stress has been associated with deleterious effects on both larger and smaller vessels. However, a set-point theory of WSS has been previously proposed (reviewed in [2]), and we argue here that the set-point is dynamic and locally controlled – thus explaining how low flow-conditions, as in our case, contributes to an increased state of health in microvessels (in comparison to maintained static culture). The number of perfused segments decreases over time in the absence of perfusion, thus resulting in increased vascular resistance and uneven distributions of fluid and nutrients.

Establishing long-term flow (beyond 48 hours) in this macro-scale device with our microvessels is on-going; however, this has been recently shown using a different system in our group [49]. We hypothesize that establishing a dynamic flow regime may be required to maintain the viability and perfusion of *in vitro* vessels, since significant remodeling and an inability to maintain vessels has been observed (96 hours). Recently, we also demonstrated that long-term (48 hour) flow induces complex-changes in cathepsin activity [50], a phenomenon largely attributed to fibroblasts. Thus, minute changes in flow have a considerable impact on whole-system integrity leading to altered extravascular matrix properties, which in turn promote vessel remodeling. Another limitation is the use of HUVEC as a source of primary endothelial cells, as they have limited proliferative capacity. Recent evidence has shown that a transient re-introduction of an ETS variant transcription factor (ETV2) in HUVEC promotes tissue-specific vessel formation and increased proliferative capacity, as demonstrated at 4 weeks post re-vascularization of rat intestines [51]. We postulate that reduced endothelial proliferation capacity (and possibly senescence) adds to the long-term reduction in vessel stability *in vitro*.

Flow-conditioning under low flow conditions (resulting in <1 Pa WSS) has a protective effect in microvessels – implying a beneficial effect of introducing flow in a timely manner to the microcirculation. Interestingly, pre-conditioning vessels with blood flow has been shown to protect against injury in larger cardiac [52] and hepatic vessels [53] *in vivo*. Yet, reperfusion of small microvessels is typically associated with acute infiltration and accumulation of leukocytes, vasoconstriction, and endothelial damage [3]. There is a clear difference in mechanosensitivity (and range) to flow conditions between larger and smaller vessels, and altered hemodynamics beyond a tolerated level can deleteriously affect vascular health. It will be important in future work to distinguish the bounds of mechanosensitivity in response to flow conditions – which will provide insight into microvessel dysfunction in systemic disease [54]. While our microvessels did not undergo true reperfusion, since static conditions were maintained until flow is applied, it is clear that flow-conditioning does maintain vascular health temporarily. Future work will involve culture of vessels under continuous flow from the outset of seeding and address this need. Overall, local dysregulation of flow patterns by regressive vessel remodelling may contribute to systemic disease progression.

## Acknowledgments

KH would like to acknowledge Jean Carlos Serrano Flores from MIT for help setting up initial interstitial flow and FRAP experiments and to Giovanni Offeddu for feedback on the manuscript. KH was partially funded by a National Science and Engineering Research Council (NSERC) postdoctoral fellowship and by the National Science Foundation (CBET-0939511).

## Methods

### Microvascular network formation under interstitial flow

Human umbilical vein endothelial cells (HUVEC) and normal human lung fibroblasts (HLF) were purchased from Lonza and cultured in EGM-2MV and FGM-2, respectively. HUVEC were transduced to express cytoplasmic RFP, as described earlier. Both cell types were used between passages 7-10. Cells were cultured until near-confluent prior to detachment and mixing of 6M HUVEC/mL and 1.2M HLF/mL (a 5 to 1 ratio) in a fibrin gel (final concentration 3mg/mL). The cell-gel mix was seeded into Aim microfluidic chips (Aim Biotech) and cultured for 5 days at 37°C in a 5% CO_2_ incubator. Using adapters and syringes, an interstitial flow (IF) was generated across the gel (see Aim Biotech protocol for details) 24hrs after initial seeding. An intermittent interstitial flow was maintained by re-adjusting the fluid volumetric drop every 4 hours for 12 hours (3 times daily) for 4 days following seeding. Several differential fluid volumes were used to examine the time in which IF ceased (^~^4hrs). Interstitial flow velocity was measured using FRAP, as previously performed [11, 55]. Briefly, a pressure difference was generated across the gel using media supplemented with FITC dextran. A circular spot of ^~^200 microns was bleached then observed to recover in a single xy-plane (moving in the direction of fluid flow). The centroid of this bleached (and then recovered) spot was tracked using Matlab (frap_analysis, based on the Hankel transform method [56]), from which velocity was estimated.

Live confocal images (Olympus IX81) were taken on day 3-5, and morphologic parameters were characterized using NIH ImageJ macros, as previously described [20]. Here, time-dependent changes are shown with respect to the relative change from the same device.

### Device fabrication & application of continuous flow

For macro-scale fluidic devices, a negative mold was generated using 1mm thick laser-cut acrylic (Astra Products). Polydimethylsiloxane (PDMS) was then used to generate devices according to the manufacturer’s recommended ratio of elastomer base to cross-linker (Ellsworth). Following air-plasma (Harrick) bonding to glass coverslips, devices were baked for a minimum of 24 hours at 60-70°C, to ensure that they returned to a hydrophobic state.

Microvessels were formed in the devices by seeding HUVEC and HLFs, as outlined above (similar to the microfluidic devices). For the macro-scale devices, the total volume of cell-gel mixture is ^~^100μL (>10x that of an AimChip). Devices were cultured in a 37°C, 5% CO_2_, incubator for 7 days prior to imaging and setting up continuous flow. At day 7, microfluidic devices were fitted with large custom media reservoirs. An in-house pump was used to drive flow continuously using a pressure gradient across the gel. Briefly, air pressure was used to displace fluid through tubing connecting the inlet and outlet sides of the media reservoirs. Pressure driven flow was maintained across the gel (and through the microvessels) by pumping fluid from the outlet to the inlet reservoir at a rate faster (^~^0.75mL/min) than intravascular fluid flow. Pressure gradients were maintained at ^~^5mm of H_2_O (50 Pa).

### Endothelial permeability measurements

Endothelial permeability to solutes was measured at day 9, either from statically cultured samples or from those exposed to continuous flow for 48hrs. Briefly, 70kDa FITC dextran was perfused into the microvascular networks by generating a slight pressure gradient across the gel of the device, following which the pressure was stabilized. Confocal images were captured at 0, 5, and 10 minutes, from which permeability measurements were made, as reported previously [20].

### Quantitative real time PCR assays

Samples were cut from the devices with a scalpel, the fibrin gel dissolved using a combination of 10% (25mg/mL) Natto Kinase (Japan Biosciences Ltd.) and 10%Accutase (Innovative Cell Technologies) in PBS, and then RNA was collected using an RNeasy Mini kit (Qiagen). Quality of RNA was confirmed using a NanoDrop (ThermoFisher). A High-Capacity RNA-to-cDNA™ Kit (ThermoFisher) was used to prepare cDNA on a BioRad thermal cycler. Changes in gene expression between microvessels exposed to flow and those cultured under static conditions were measured using a GeneQuery™ Human Shear Stress and Mechanotransduction qPCR Array Kit (ScienCell). The qPCR cycling protocol was performed on a Roche 96 LightCycler. All 5 housekeeping genes (β-actin, GAPDH, LDHA, NONO, PPIH) were used to normalized the data using the delta-delta-Cq method as outlined in the GeneQuery array protocol.

For genes of interest, TaqMan assays: Hs01110250_m1 (HMOX1), Hs01003372_m1 (VCAM1), Hs00892478_g1 (GPC1), Hs01080223_m1 (BBC3), and Hs00153153_m1 (HIF1A) were used along with Hs01060665_g1 (ACTB) as a control. All assays were performed using TaqMan Fast Advanced Master mix (ThermoFisher) on a 7900HT Fast Real-time PCR system (Applied Biosystems) according to recommended protocols.

### Nitric oxide, ROS, and caspase 3/7 detection

NO detection was performed by incubating microvessels with 5μM DAF-2 (Calbiochem) for 30 minutes at 37°C prior to imaging at 2-minute intervals. Samples were first imaged without flow, and then under flow by imparting an immediate pressure difference of 2 mm H_2_O (using syringes). Reactive oxygen species were detected with ROS-ID^®^ Hypoxia/Oxidative stress detection kit (Enzo Life Sciences) according to manufacturer’s protocol. Apoptosis was detected by incubating devices with 5μM of CellEvent™Caspase 3/7 Green detection reagent (Invitrogen) for 30 minutes at 37°C.

### Immunocytochemistry staining & ELISA

For ICC, samples were fixed in 4% PFA, rinsed with PBS, permeabilized (0.1%TritonX in PBS), and then incubated in blocking buffer (PBS + 5% BSA + 3% goat serum) for several hours, prior to incubation with primary antibodies overnight at 4°C. The following day, devices were rinsed with wash buffer (0.5% BSA in PBS) thoroughly prior to incubation for several hours with the corresponding conjugated antibody.

Supernatants were collected from devices at day 9 following culture under static conditions or 48 hours of flow (days 7-9). ELISA was performed according to manufacturer’s protocols for Ang-1 (ThermoFisher) and Ang-2 (Invitrogen) detection.

### Computational fluid dynamics modeling of microvascular perfusion

Vessel geometries were reconstructed from confocal images (512×512 pixels at a resolution of 2.485 μm/pixel) and slices (z-direction) of 5 μm. The network was captured with a 4×2 grid of images along x-y directions to cover large region of the device. Images were stitched and stacked along the z-direction by means of in-house software developed in Matlab (The MathWorks, Natick, Massachusetts, USA). A double thresholding algorithm was applied on the 3D image volume to isolate the region of interest of the vessels, from which boundaries were extracted with a marching cube algorithm to compute the 3D iso-surface defining the vessels walls (as in [23]). The final 3D geometry was obtained in Meshmixer (Autodesk, San Rafael, CA, USA) by performing cuts on the y-z plane to define inlet and outlet closed surfaces. The final geometry was processed to compute an estimate of 3D porosity, as the ratio between the enclosed vessels volume and the stacked-images volume, and of the vessel diameters on y-z cross-sectional planes.

Numerical modeling of vessels perfusion was performed by importing the 3D vessel geometry into Ansys ICEM CFD (Ansys, Canonsburg, PA, USA) to define the discretized numerical grid, consisting of approximately 5.5M tetrahedral elements. Boundary layers were imposed to improve flow gradient estimation near the vessel walls. Modeling of vessel perfusion was accomplished using the numerical solver Ansys Fluent (Ansys, Canonsburg, PA, USA). Computational fluid dynamics simulations were performed considering a laminar flow condition and no slip at the vessel wall. The flowing fluid was assumed to be Newtonian with a dynamic viscosity of 9.4×10^−4^Kg·m^−1^·s^−1^ and density of 998.2 kg·m^−3^ working at 37°C (values reported for DMEM +10%FBS) [24]. A static pressure drop of 1cm H_2_0 was imposed between inlet and outlets (across the entirety of vessels), while the lateral boundaries were treated as periodic surfaces to ensure continuity of the pressure/velocity in the fluid grid. This simulation was considered (2x) representative of the standard experimental set-up; additional pressure-drops (7mm H_2_O, 13mm H_2_O) were tested to verify linearity of hemodynamic variables variation as well as expected velocity outputs (Figure S5). Simulations results were exported and post-processed in Matlab to compute the distribution within the vessels network of flow velocity, shear rate, WSS and vorticity values. Sensitivity analyses and grid convergence tests were run to ensure the accuracy of performed numerical simulations. Streamlines were computed and visualized to assess, qualitatively, the evolution of flow to be consistent with computed simulations.

### Statistics

Unless noted otherwise, values shown represent mean ± standard error of the mean (s.e.m.), and student’s t-test with P<0.05 indicating significant differences between means. Where significant differences between multiple samples are tested, one-way ANOVA with post-hoc Tukey test is used for means comparison.

