## Supplementary Material for "Physiologic flow-conditioning limits vascular dysfunction in engineered human capillaries"

### Supplementary Figures

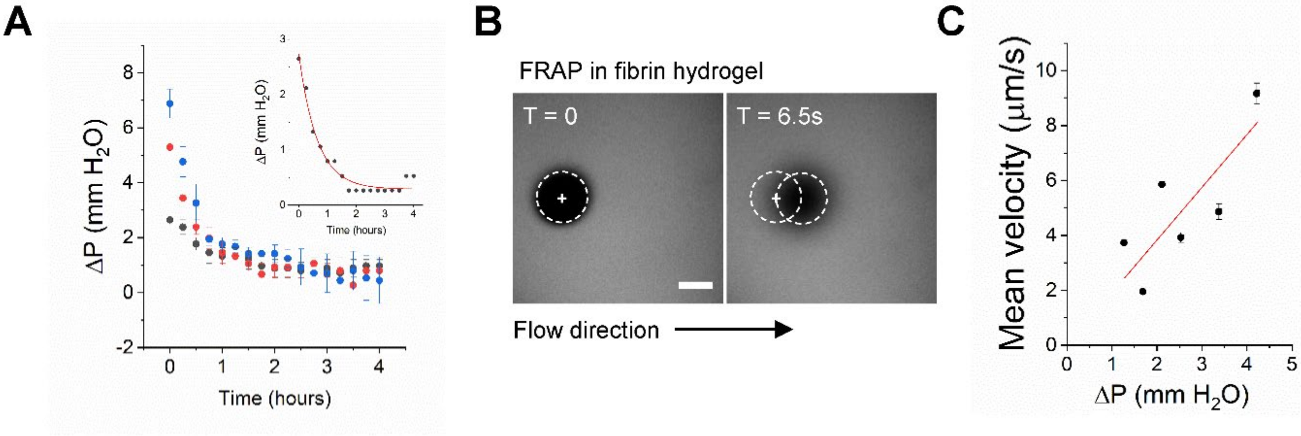


**Figure S1**: Characterization of flow velocity in fibrin hydrogels. A) Volume change of media measured over time for AimChip IF experiments. Black ≈ 2.5, red ≈ 5, and blue ≈ 10 are mm$H_2O$ initial pressure differences. Inset is an example of exponential decay fit ($R^2$=0.967) of one measurement with starting pressure difference set at 2.5mm $H_2O$. B) FRAP measurements were made in large macro-fluidic chips containing fibrin. A circular ROI was bleached and consecutive images were used to track the flow velocity for a given volume difference. Dashed circles indicate the bleached region in a FITC dextran-filled gel. C) Mean velocity is shown for different fluid volume change (pressure difference across the gel). A linear relationship exists between the volume difference and fluid flow velocity in the gel.

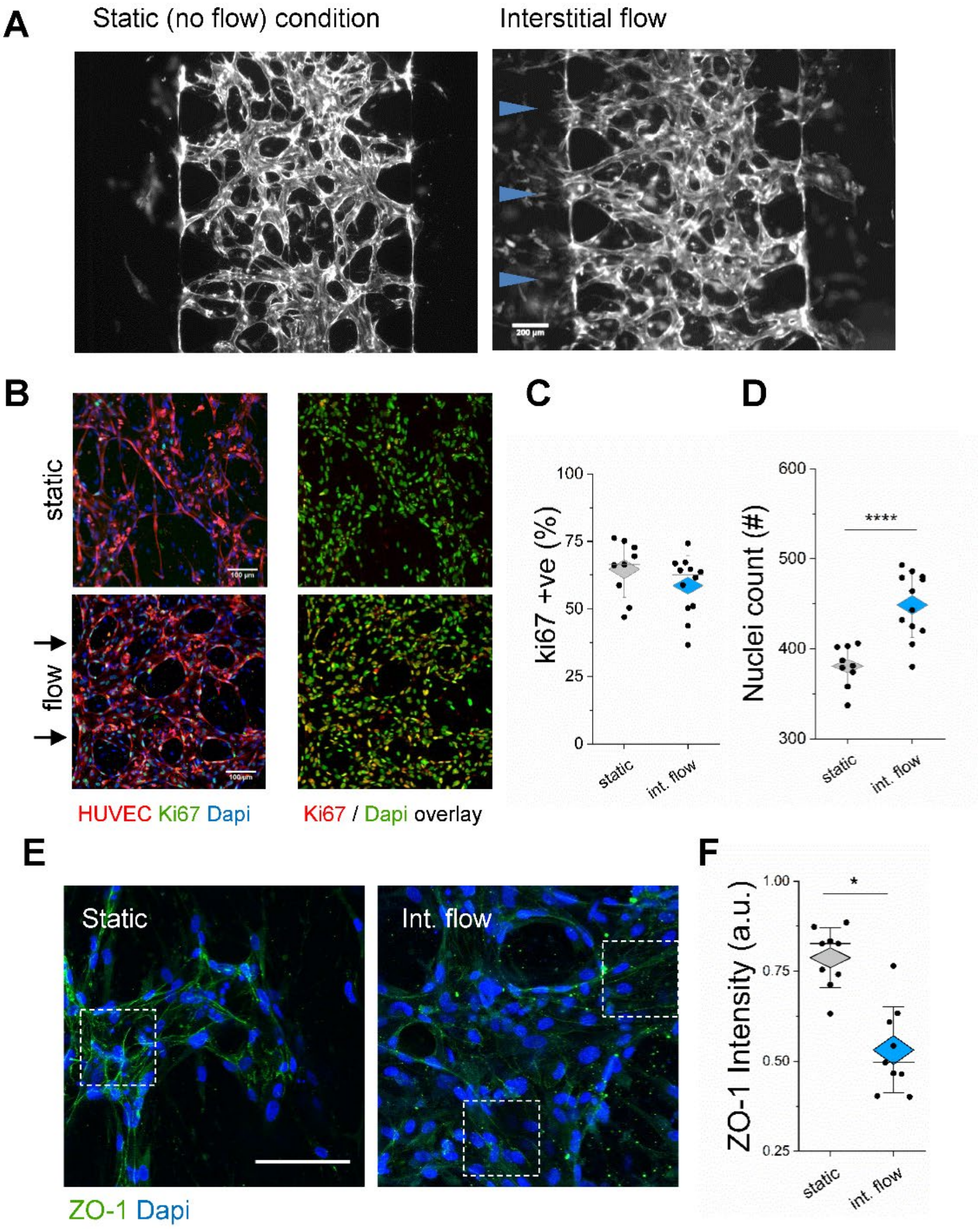


**Figure S2**: Interstitial flow increases vessel formation and alters endothelial junction stability. A) Epi-fluorescent images demonstrate significant morphological differences between vessels (grey) grown under static or interstitial flow conditions at day 4. Blue arrows indicated flow direction in right image. B) Example of fixed and stained samples from each condition demonstrating proliferation of HUVEC (red) and fibroblasts (non-fluorescent) in the system (left panel), with a red/green overlay of ki67+ nuclei (red) and all nuclei (blue) on the right panel. C) Corresponding measurements of ki67+ nuclei were made from stained images, such as those in (B). D) Nuclei count between conditions was significantly different, despite seeding with the same cell density on day 0. E) Fixed and stained images of tight-junction protein ZO-1. Regions of interest are highlighted by dashed box. Intensity was measured in multiple ROIs and normalized to nuclei (DAPI) intensity for both static and flow conditions. Scale bar is 100µm. F) ZO-1 showed increased expression for the case of vessels growing under static conditions as opposed to flow, as measured from fixed/stained images, as in (E). Significance is indicated by unpaired t-tests between static and flow, with *P<0.05, ****P<0.0001.

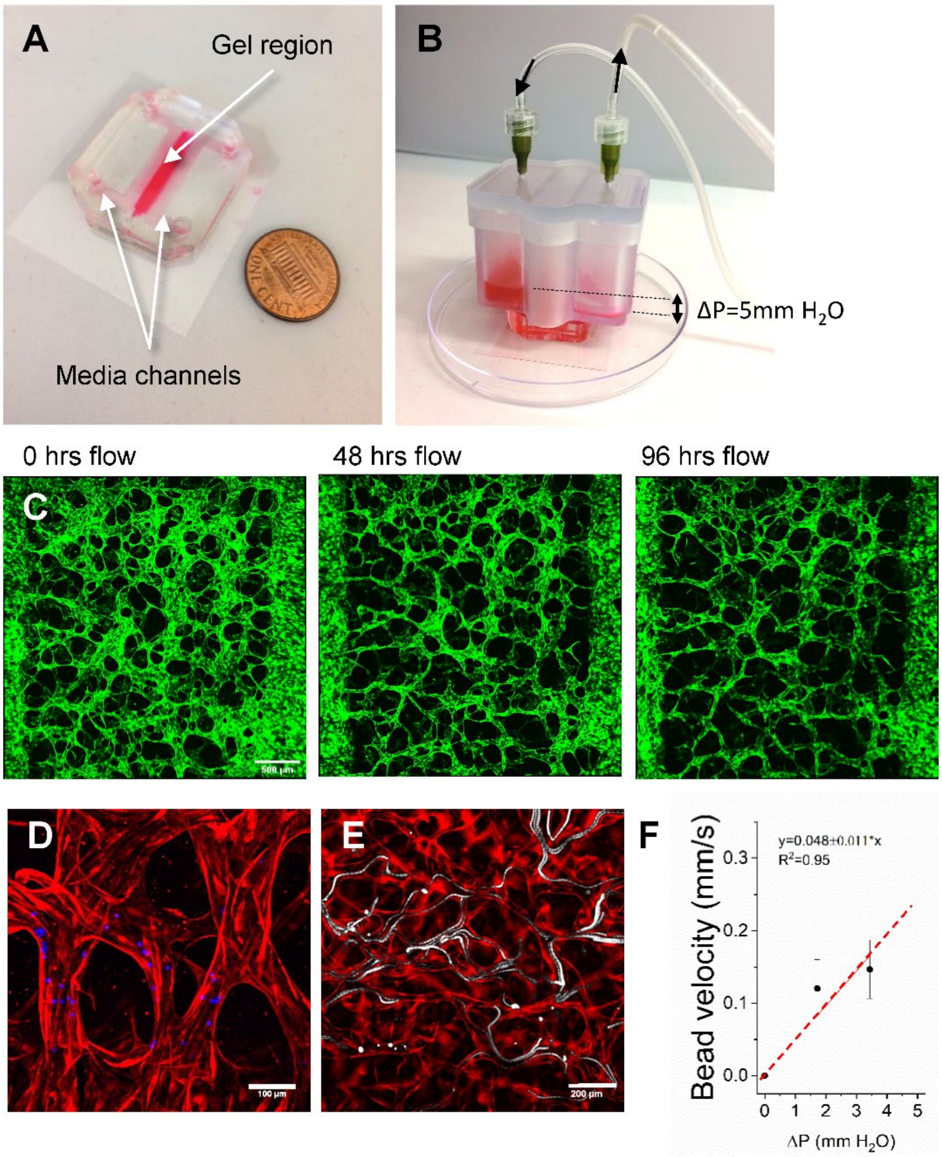


**Figure S3**: Set-up of microfluidic device and flow chamber. A) A macro-scale device is shown filled with PBS + food colorant. The dark pink region highlights the gel containing region (fibrin hydrogel). B) Image showing the large media reservoir for generating a pressure gradient across a macro-scale device (as shown in A). Fluid is re-circulated using a solenoid and air-pressure controlled pump. C) Large merged region of vessels (same sample) showing vessels rarefaction despite continuous flow for 96 hours. Scale is 500 microns. D) Z-projection demonstrating 10 micron beads (blue) inside luminal vessels (red). E) Epi-fluorescent image demonstrating a time-trace of 10 micron beads (white) flowing through the hollow vessels (HUVEC-RFP). F) Mean bead velocity as measured across several devices for different pressure gradients. Pressure gradients correspond to 200µl and 400µL volume difference (with 5mL syringes). Average bead velocities are shown from 200-750 beads analyzed each.

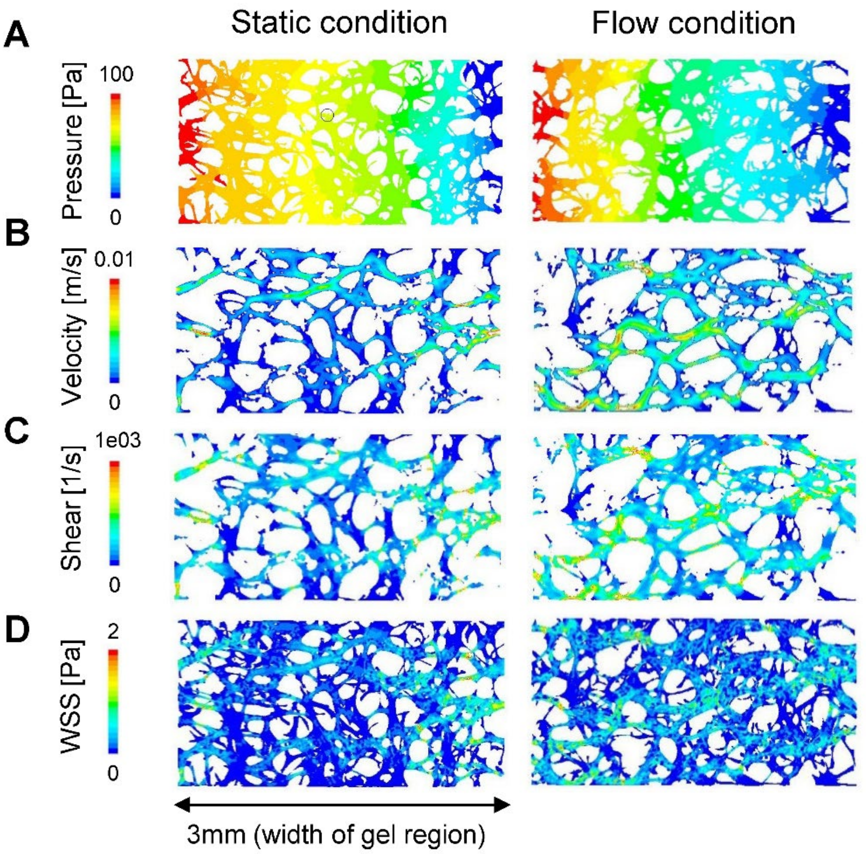


**Figure S4**: Computational fluid dynamic models of vessels treated to static and continuous flow conditions. A-D) A comparison of one example each of vessels cultured under static versus flow conditions for 48hrs (day 7-9 in culture). Confocal images were converted to 3D surfaces and used in CFD modeling (n=3 each). A) A pressure gradient was simulated in the CFD to mimic similar conditions to those used in vitro. Under a 10mm $H_2O$ pressure gradient (~100 Pa), local differences are shown between static and flow-conditioned vessels, the latter of which is more homogenously perfusable. Shown are example distributions of B) mean velocity, C) shear rate, and D) WSS, for the two samples.

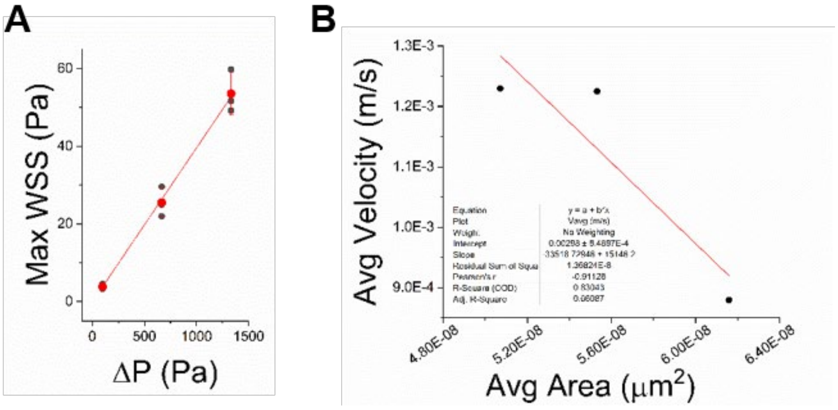


**Figure S5**: Validation of CFD simulations. A) Increasing the pressure gradient across the vessels results in a linear ($R^2$=0.999) increase in WSS, as expected. B) Average fluid velocity increases with decreasing vessel area, with volumetric flow unchanged (as expected).
